# Myelin antigen-specific effector CD8^+^ T cells induce chronic CNS autoimmunity in a CD4^+^ T cell-dependent manner

**DOI:** 10.1101/2025.03.25.645315

**Authors:** Irshad Akbar, Prenitha Mercy Ignatius Arokia Doss, Joanie Baillargeon, Mohamed Reda Fazazi, Ana C Anderson, Hans Lassmann, Manu Rangachari

## Abstract

Both CD4^+^ and CD8^+^ T cells play critical roles in the immunopathogenesis of multiple sclerosis (MS). 1C6 T cell receptor transgenic (TcR-Tg) mice on the nonobese diabetic (NOD) background have a MOG_[35-55]-_specific, MHC class II-restricted, TcR that selects for both CD4^+^ and CD8^+^ T cells. Adoptive transfer of 1C6 CD4^+^ T helper 17 (Th17) cells can induce experimental autoimmune encephalomyelitis (EAE) with a progressive disease course. In the current study, we assessed the function and pathogenicity of 1C6 CD8^+^ T cells. We found that they proliferated and produced inflammatory cytokines in response to MOG_[35-55]_ peptide under both T cytotoxic 1 (Tc1) and Tc17 differentiation conditions, albeit with reduced expansion relative to their Th1 or Th17 counterparts. Both 1C6 Tc1 and Tc17 cells were able to induce EAE upon adoptive transfer to NOD.*Scid* mice. Intriguingly, we noted *in vivo* expansion of CD4^+^ T cells in the spleen and CNS of NOD.*Scid* recipients as well as in lymphocyte-sufficient animals, despite 1C6 Tc cells being purified on CD8 expression prior to transfer. Furthermore, 1C6 Tc17 cells expressed ThPOK, a master differentiation factor for CD4^+^ T cells. Finally, anti-CD4^+^ T cell blockade abrogated CD8^+^ T cell infiltration of the CNS and disease induction in Tc17 recipient mice. Our data provide insight into the interplay of CD4^+^ and CD8^+^ T cells in CNS autoimmunity.

## Introduction

Multiple sclerosis (MS) is an autoimmune and neurodegenerative disease of the central nervous system (CNS), affecting nearly 2 million people worldwide (1), that leads to demyelination and neurodegeneration in the CNS (2). It is suggested to be mediated by an adaptive immune cell-mediated attack against CNS myelin constituents, though reactivity to other CNS antigens or to Epstein-Barr virus (EBV) has also been observed (3, 4). Approximately 80% of people living with MS (pwMS) will initially experience a relapse-onset form of the disease that is relatively controllable by existing disease-modifying treatments. However, 30-60% of these individuals will eventually transition to secondary-progressive (SP) MS, while 20% of all pwMS show progressive disease from onset (primary progressive, PP). Treatment for progressive forms of MS is currently limited, leaving these individuals to face decades of accumulating symptoms (5, 6).

MS risk is strongly associated with polymorphisms in HLA class II genes (7), indicating the involvement of pathogenic CD4^+^ T cells. Further, animal models of CNS autoimmunity have delineated a crucial role for IFNγ-producing CD4^+^ T helper 1 (Th1) and IL-17-producing Th17 responses (8). However, CD8^+^ T cells actually outnumber CD4^+^ T cells in MS lesions (9, 10). Like CD4^+^ T cells, effector CD8^+^ T cells can differentiate into IFNγ-skewed (T cytotoxic 1; Tc1) or IL-17-positive (Tc17) subtypes (11). Tc17 cells in particular have been detected in CNS lesions in both MS (12, 13) and neuromyelitis optica spectrum disorder (14); however, the role(s) played by these CD8^+^ cell subsets in autoimmune CNS injury are incompletely understood.

Experimental autoimmune encephalomyelitis (EAE) is an animal model that permits the study of numerous elements of CNS autoimmunity. EAE can be induced either by active immunization with encephalitogenic peptides or proteins, or by the transfer of CNS-antigen specific lymphocytes. Different models of EAE exhibit variations in clinical presentation, depending on factors such as method of disease induction, target immunogen, and/or genetic background (8). The majority of EAE models feature a CD4^+^ T cell-initiated pathology, either because they depend on immunization with MHC class II-restricted autoantigenic peptide or because they require the adoptive transfer of CD4^+^ T cells. Nevertheless, effector CD8^+^ cells accumulate in murine models of MS (13, 15–17) and a number of CD8^+^ T cell-dependent EAE models have been described; together, they show that class I-restricted CD8^+^ T cell responses that are characterized by the influx of inflammatory CD8^+^ T cells can infiltrate the CNS and cause inflammatory lesions in both white and grey matter (18–22).

Nonobese diabetic (NOD) background mice can develop a relapsing/chronic disease pattern upon active immunization with myelin oligodendrocyte glycoprotein 35-55 (MOG_[35-55]_) peptide. NOD-EAE has thus been used as a model of the immune pathogenesis of SPMS (23–26). NOD-background T cell receptor transgenic (TcR-Tg) 1C6 mice, possess an antigenic specificity for MOG_[35-55]_ (27). Adoptive transfer of either Th1 or Th17 1C6 cells to NOD.*Scid* hosts induces EAE; in particular, the transfer of male 1C6 Th17 cells elicits a severe progressive clinical course (28). Surprisingly, despite the fact that the 1C6 TcR is MHC class II-restricted, both CD4^+^ and CD8^+^ MOG_[35-55]_-reactive T cells are detected in the immune periphery of these animals (27). The goal of the current study was thus to investigate the function of 1C6 Tc1 and Tc17 cells, and to ascertain their capacity to induce EAE upon adoptive transfer.

## Results

### 1C6 CD8^+^ T cells respond suboptimally to MOG_[35-55]_ but show no intrinsic proliferative defect

The MHC class II-restricted 1C6 TcR is expressed by both mature CD8^+^ and CD4^+^ T cells, even though CD8^+^ T cells typically possess a TcR that is MHC class I-restricted. It was previously shown that 1C6 CD8^+^ T cells proliferate in response to MOG_[35-55]_ and that this is abrogated upon blockade of the NOD class II allele I-A^g7^ (27). Here, we directly compared the proliferative capacity of 1C6 Tc1 and Tc17 cells to that of Th1 and Th17 cells. We first purified naïve CD62L^hi^ populations (29) of CD4^+^ and CD8^+^ T cells from the peripheral lymphoid tissues of 1C6 mice and cultured them under well-defined type-1 or type-17 culture conditions (30) in the presence of MOG_[35-55]_-pulsed irradiated splenocytes. Under these conditions, both CD8^+^ Tc1 and Tc17 cultures showed reduced cellularity relative to their CD4^+^ Th1 or Th17 counterparts (Figure 1A). Interestingly, however, the proliferative index of Tc1 cultures was higher than that of Th1 comparators, while no differences between Tc17 and Th17 cells were observed. Thus, 1C6 Tc1 and Tc17 cultures did not expand as well as Th1/Th17 cells upon stimulation with MOG_[35-55]_; however, when considering only cells that had undergone at least one division, 1C6 Tc1 and Tc17 cells proliferated as well as their respective effector CD4^+^ T cell counterparts. We next assessed proliferative responses of 1C6 CD8^+^ Tc1 and Tc17 cells versus CD4^+^ Th1 and Th17 cells upon stimulation with plate-bound agonistic antibodies to CD3 and CD28 (Figure 1B). We found no differences in either cellularity or proliferative index between Tc1 and Th1 cells or between Tc17 and Th17 cells under these conditions, which trigger T cell responsiveness in an antigen-independent manner. Overall, our data indicated that 1C6 Tc1 and Tc17 cells responded to cognate MOG_[35-55]_ antigen, though they expanded less capably than Th1 and Th17 cells. Further, 1C6 effector CD8^+^ T cells show no intrinsic proliferative defect.

**Figure 1.**
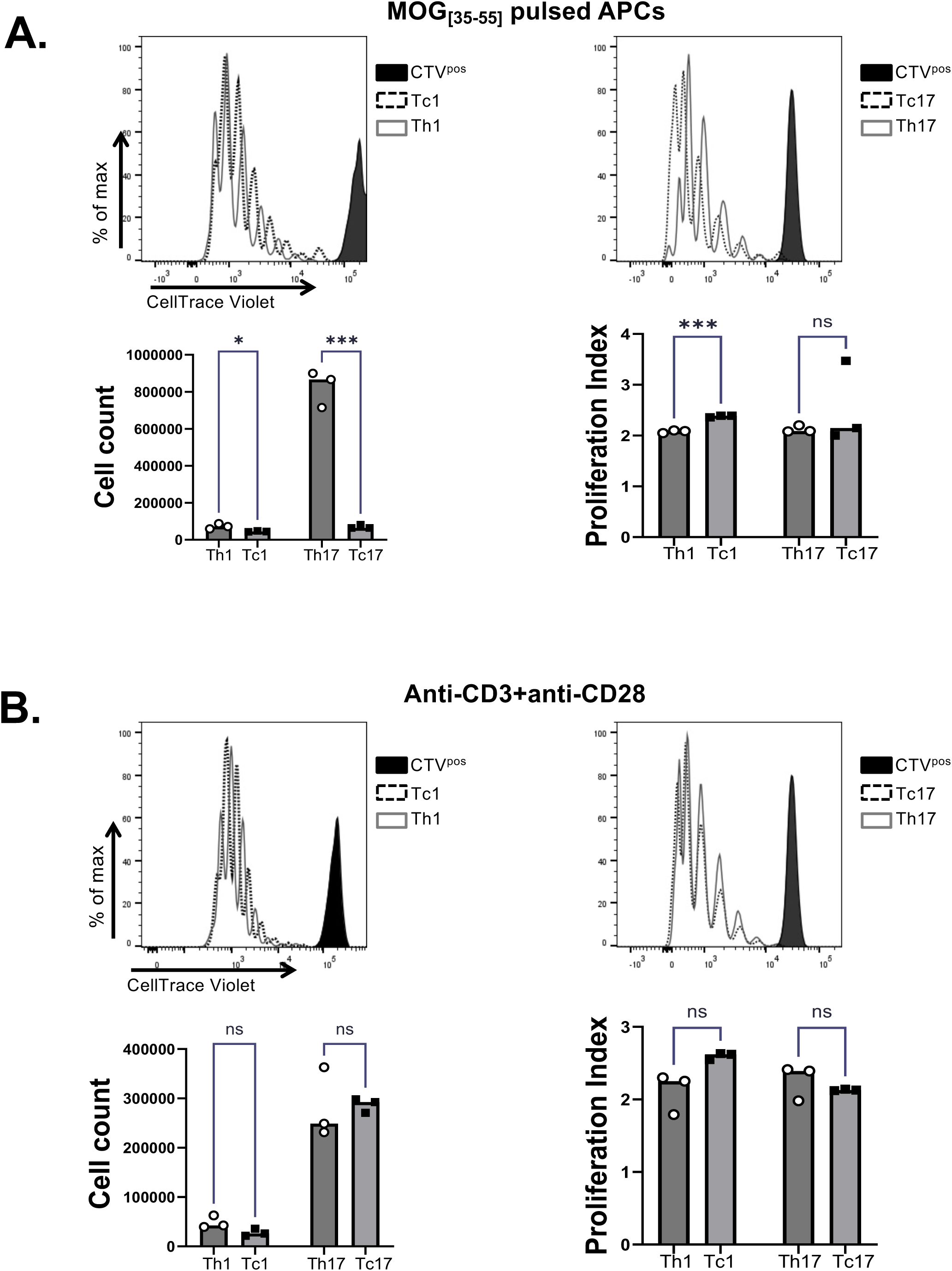
1C6 CD8^+^ T cells expand suboptimally to MOG_[35-55]_ peptide but show no intrinsic proliferative defect. Female CD4^+^CD62L^hi^ 1C6 CD4^+^ and CD8^+^ T cells were stimulated with MOG_[35-55]_-pulsed APCs (**A**) or with plate-bound anti-CD3 + anti-CD28 (**B**) under type-1 (Th1, Tc1) or type-17 (Th17, Tc17) differentiation conditions. Cell proliferation was assessed by CellTrace Violet dilution after 3 days of culture. *, p<0.05; ***, p<0.001; n.s., not significant; two-tailed *t*-test. Each replicate represents a culture derived from a distinct mouse. CTV^pos^: CellTrace Violet-stained, unstimulated cells. Data representative of 3 experiments.

### Assessment of cytokine production from 1C6 effector T cells

We next examined inflammatory cytokine production from 1C6 effector Tc and Th cells. Upon stimulation with MOG_[35-55]_-pulsed splenocytes, a lower frequency of Tc1 cells produced IFNγ and TNF compared to Th1 counterparts (Fig. 2A). Under type-17 conditions, a greater frequency of MOG_[35-55]_-stimulated Tc17 cells produced IFNγ than Th17 comparators (Fig. 2A), likely reflecting the propensity of CD8^+^ T cells to generate IFNγ. Further, we observed an increased frequency of TNF-producing 1C6 CD8^+^ T cells among Tc17 versus Th17 cells. There were no differences in the frequency of IL-17-producing cells. Under plate-bound conditions (Fig. 2B), we observed increased IFNγ frequency from both Tc1 and Tc17 cells relative to Th1 and Th17 comparators. No differences in the frequency of TNF-producing cells were observed; however, 1C6 Tc17 cultures had an increased frequency of IL-17-producing cells as compared to Th17 cells. Overall, these data suggested that 1C6 Tc1 and Tc17 capably produce inflammatory cytokines.

**Figure 2.**
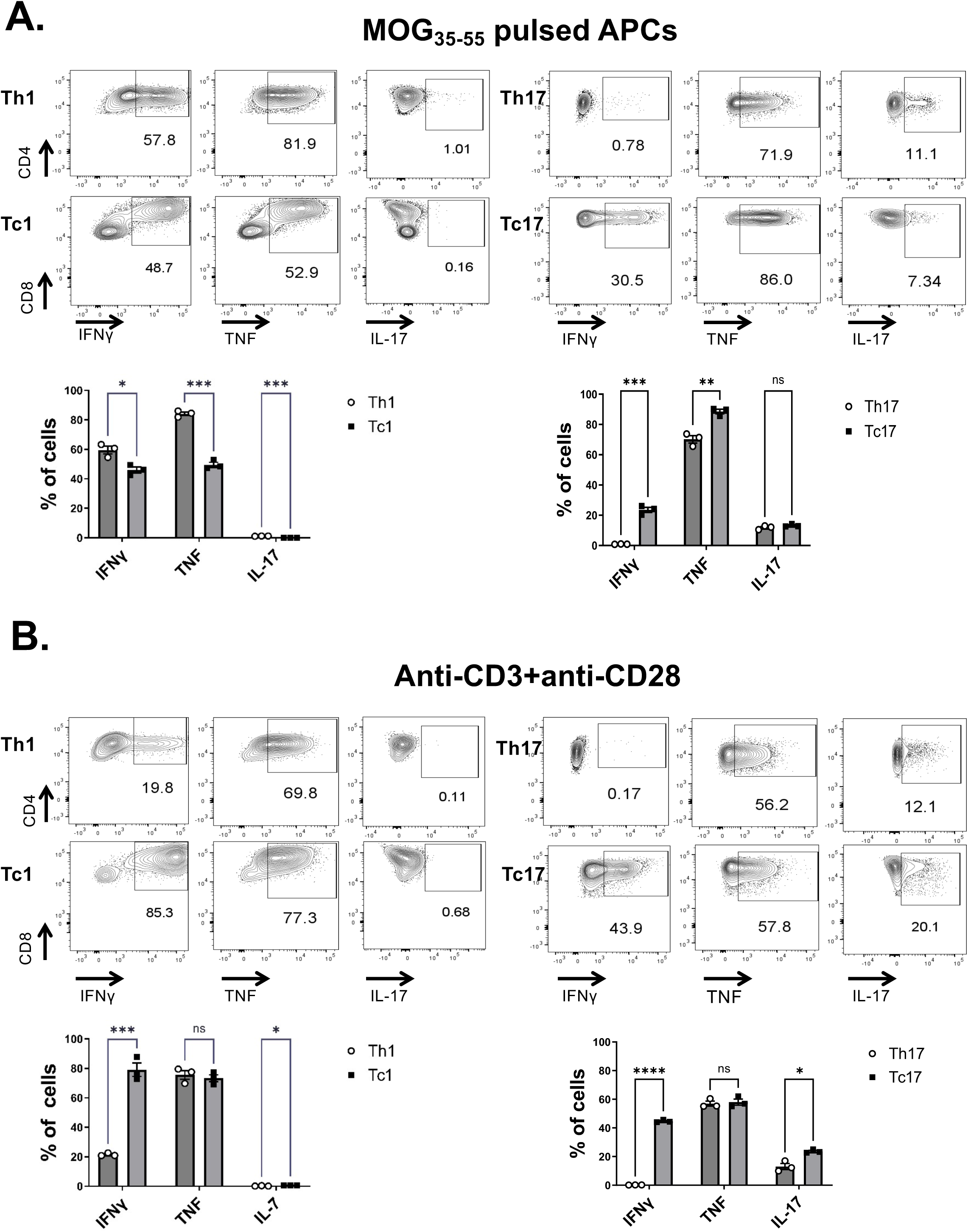
Assessment of cytokine production from 1C6 effector T cells. **A.** Female 1C6 CD4^+^ Th1 and Th17 cells, and CD8^+^ Tc1 and Tc17 cells were differentiated and stimulated with either MOG_[35-55]_-pulsed APCs (**A**) or with plate-bound anti-CD3+CD28 (**B**). Expression of the indicated cytokines was assessed by flow cytometry after 3 days of culture. Gated on live CD4^+^ or live CD8^+^ T cells. *, p<0.05; **, p<0.01; ***, p<0.001; ****, p<0.0001; n.s., not significant; two-tailed *t*-test. Each replicate represents a culture derived from a distinct mouse. Data representative of 3 experiments.

### 1C6 Tc1 and Tc17 cells adoptively transfer EAE

We next wanted to ascertain whether 1C6 effector CD8^+^ T cells were encephalitogenic. We had previously shown that plate-bound-stimulated (anti-CD3 + anti-CD28) 1C6 Th1 and Th17 cells could induce EAE upon adoptive transfer to NOD.*Scid* recipient mice (28). As we had found that 1C6 Tc1 and Tc17 cells showed robust activation *in vitro* upon the same stimulation parameters, we modified our previous experimental paradigm (28) to assess their capacity to cause EAE (Fig. 3A). We first generated female 1C6 Tc1 cells and adoptively transferred them (5×10^6^ per mouse) to female NOD.*Scid* recipients that we monitored for EAE symptoms over 70 days. In parallel, we monitored recipients of an identical number of female 1C6 Th1 cells, as well as mice that received both Th1 and Tc1 cells (2.5×10^6^ each). Tc1 recipients developed EAE of delayed onset relative to those that received either Th1 cells alone or Th1+Tc1 cells together (Fig 3B); however, their disease burden was no less than that observed in the other groups. Notably, mice that received Th1+Tc1 cells developed EAE with similar onset and disease burden compared to mice that received Th1 cells alone.

**Figure 3.**
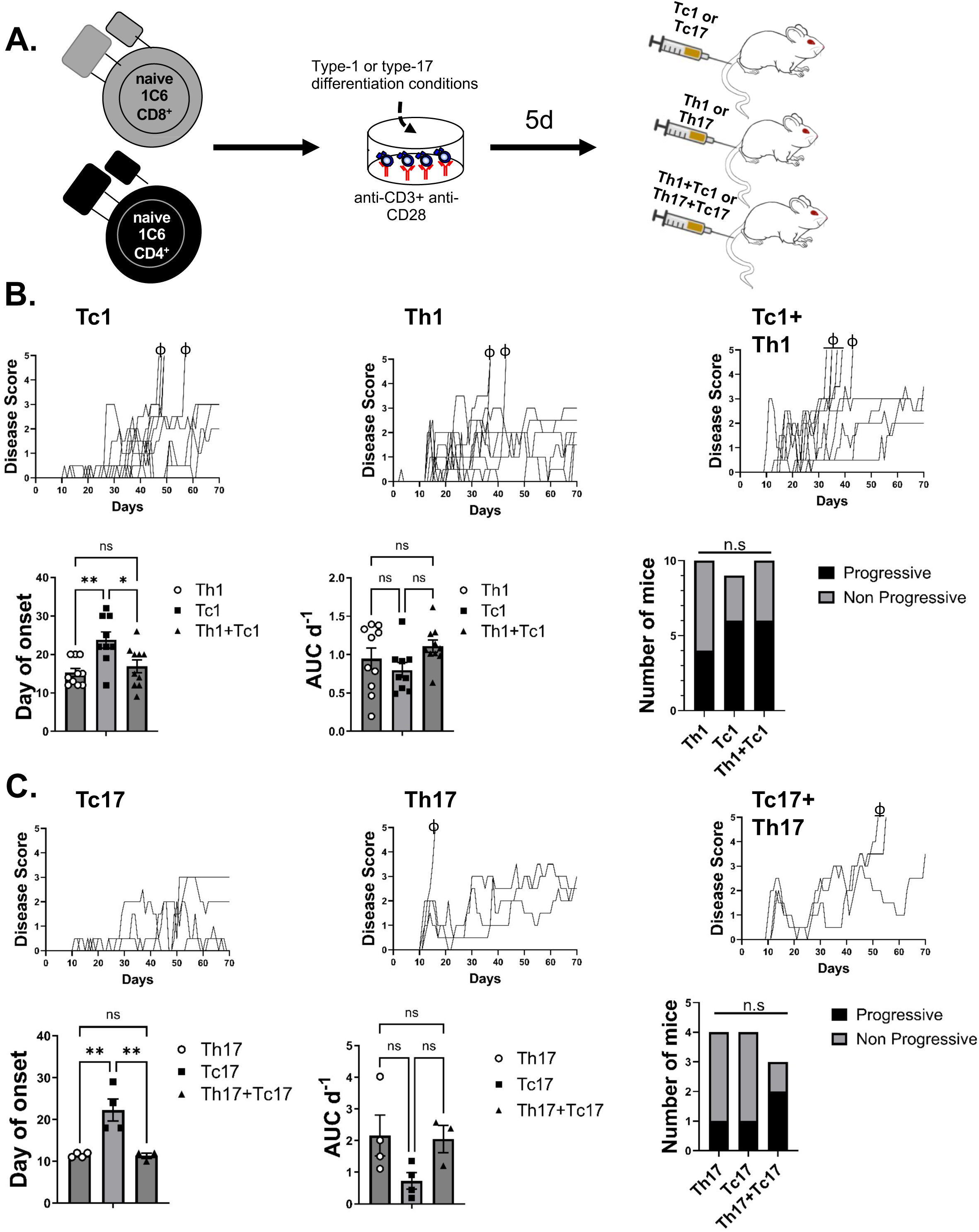
1C6 Tc1 and Tc17 cells adoptively transfer EAE. **A.** Schematic depicting experimental approach used in this Figure. **B.** Female 1C6 Tc1 and Th1 cells were differentiated in vitro and were injected i.v. into three cohorts of NOD.*Scid* mice: Tc1 alone (5×10^6^ cells, n=9), Th1 alone (5×10^6^ cells, n=10), Th1+Tc1 (2.5×10^6^ cells each, n=10). Recipients were monitored daily for signs of EAE until d70. **C**. Female 1C6 Tc17 and Th17 cells were injected i.v. into Tc17 alone (n=4), Th17 alone (n=4) or Th17+Tc17 recipients (n=3). Mice were monitored daily as in (**B**). Each symbol represents an observation from a distinct mouse. AUC d^-1^, area under curve per day. ϕ, mouse attained ethical endpoint. *, p<0.05; **, p<0.01; n.s, not significant; Tukey’s post-hoc test.

NOD-background mice are prone to developing a chronic progressive disease course (23–26). In our 1C6 adoptive transfer model, we had previously found that a modest frequency of female Th1 cell recipients also display this phenotype (28). Here, we found that 3/10 Th1 recipients displayed chronic disease, as did 5/9 Tc1 and 6/10 Th1+Tc1 recipients (Fig 3B), indicating that the transfer of Tc1 cells could also elicit a progressive phenotype in some recipient animals.

We next compared disease caused by female Tc17 cells transferred alone, female Th17 cells transferred alone, or by the co-transfer of Th17+Tc17 cells. We found that transfer of Tc17 cells alone caused disease that was of delayed onset, yet of similar overall severity, as compared to Th17 cells alone or of Th17 cells with Tc17 cells. Again, co-transfer of Th17+Tc17 cells together yielded disease of similar onset and severity as that caused by Th17 cells alone (Fig 3C). In line with our previous findings, a minority (1/4) of female Th17 cell recipients developed progressive disease; of Tc17 recipients, 1/4 showed this phenotype, as did 2/3 Th17+Tc17 recipients (Fig 3C).

We next assessed histopathological tissue damage in the CNS tissue of Tc recipient mice (3 Tc1, 1 Tc17) at endpoints. In the spinal cord, inflammation was observed in all mice studied, with demyelinating lesions seen in 3 of 4 spinal cords (Supplementary Table 1). Such lesions showed evidence of axonal loss (Figure 4A). We had previously found that the adoptive transfer of 1C6 Th cells elicits inflammatory lesions in the cerebellum, midbrain and thalamus, with demyelinating lesions seen only in the medulla (28). We therefore further characterized the lesions in the brains of Tc recipient mice. In Tc1 recipients, we observed lesions of variable size that were both inflammatory and demyelinating in nature (Supplementary Table 1). Notably, one animal presented large demyelinating lesions, with clear evidence of axonal injury, in the midbrain and thalamus (Figure 4B). The latter was of particular interest as thalamic injury is seen in the great majority of pwMS studied at either autopsy (31) or by MRI (32). Overall, our data indicated that adoptive transfer of 1C6 effector CD8^+^ Tc1 or Tc17 cells induced EAE of comparable severity to effector CD4^+^ Th1 or Th17 cells, albeit with delayed onset. Further, 1C6 Tc cells elicited pathogenic lesions in the spinal cord and brain.

**Figure 4.**
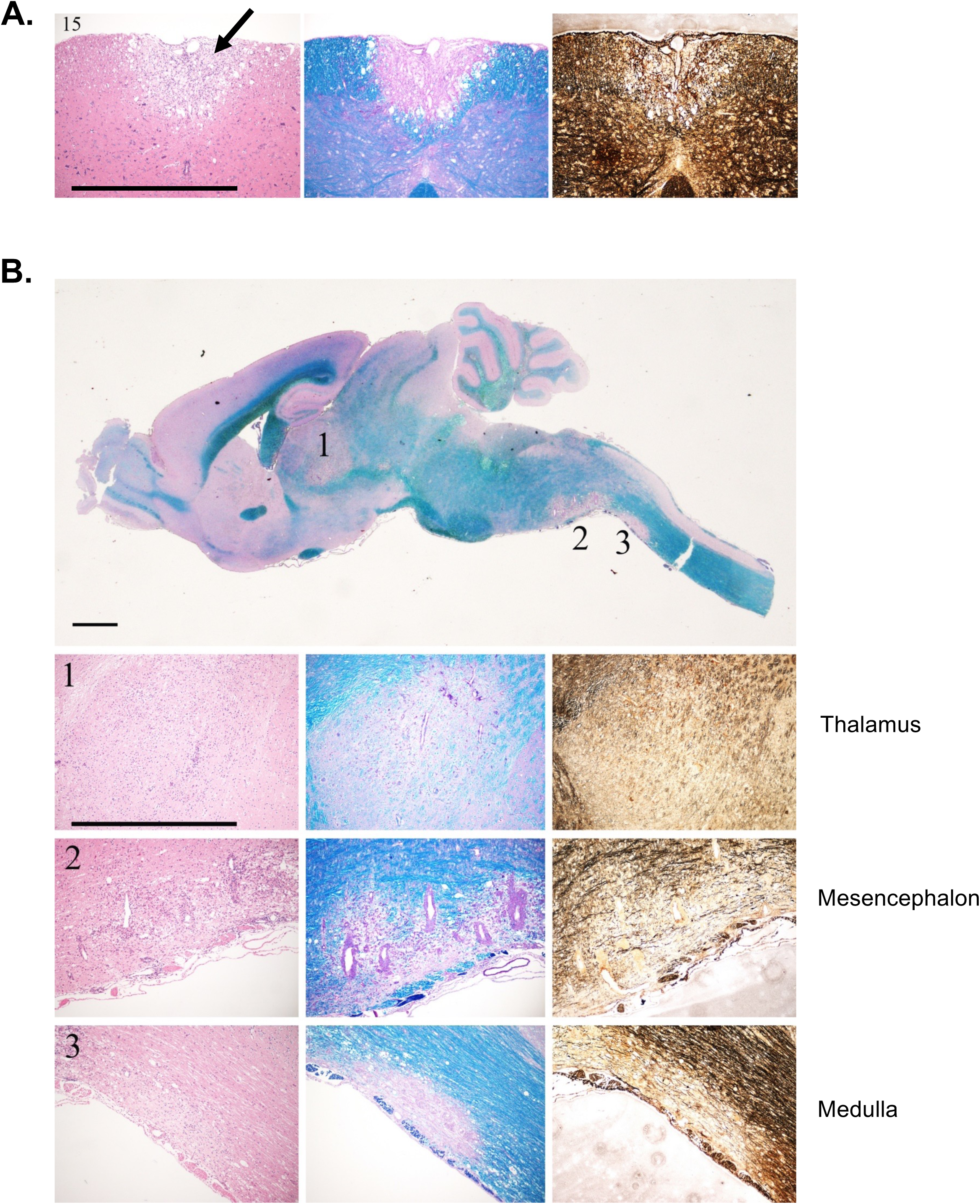
Histopathological assessment of 1C6 Tc recipients. **A.** Representative image of paraffinized and stained spinal cord tissue from a female Tc1 recipient at endpoint. *Left*, leukocytic infiltration (H&E), *middle*, demyelination (Luxol fast blue), *right*, axon damage (Bielschowsky’s stain). Arrow indicates lesion. **B.** Representative image of paraffinized and stained brain tissue from a female Tc1 recipient at endpoint with lesions in the thalamus (*top*), mesencephalon (*middle*) and medulla (*bottom*). All magnification bars represent 1mm. *Left*, leukocytic infiltration (H&E), *middle*, demyelination (Luxol fast blue), *right*, neuron damage (Bielschowsky’s stain). Related to Supplementary Table 1.

### Male and female 1C6 Tc17 cells induce EAE of similar onset and severity

To further study the effect of Tc17 cells in vivo, we adapted our culture protocol by subjecting Tc17 cells to a second 5-day round of *in vitro* differentiation (Figure 5A). This modified protocol both increased cellular yield (Figure 5B) while augmenting the encephalitogenic capacity of the cultures (Figure 5C). Thus, from this point forward, we assessed Tc-driven EAE using 1C6 CD8^+^ T cells that had been stimulated over 2 rounds prior to adoptive transfer.

**Figure 5.**
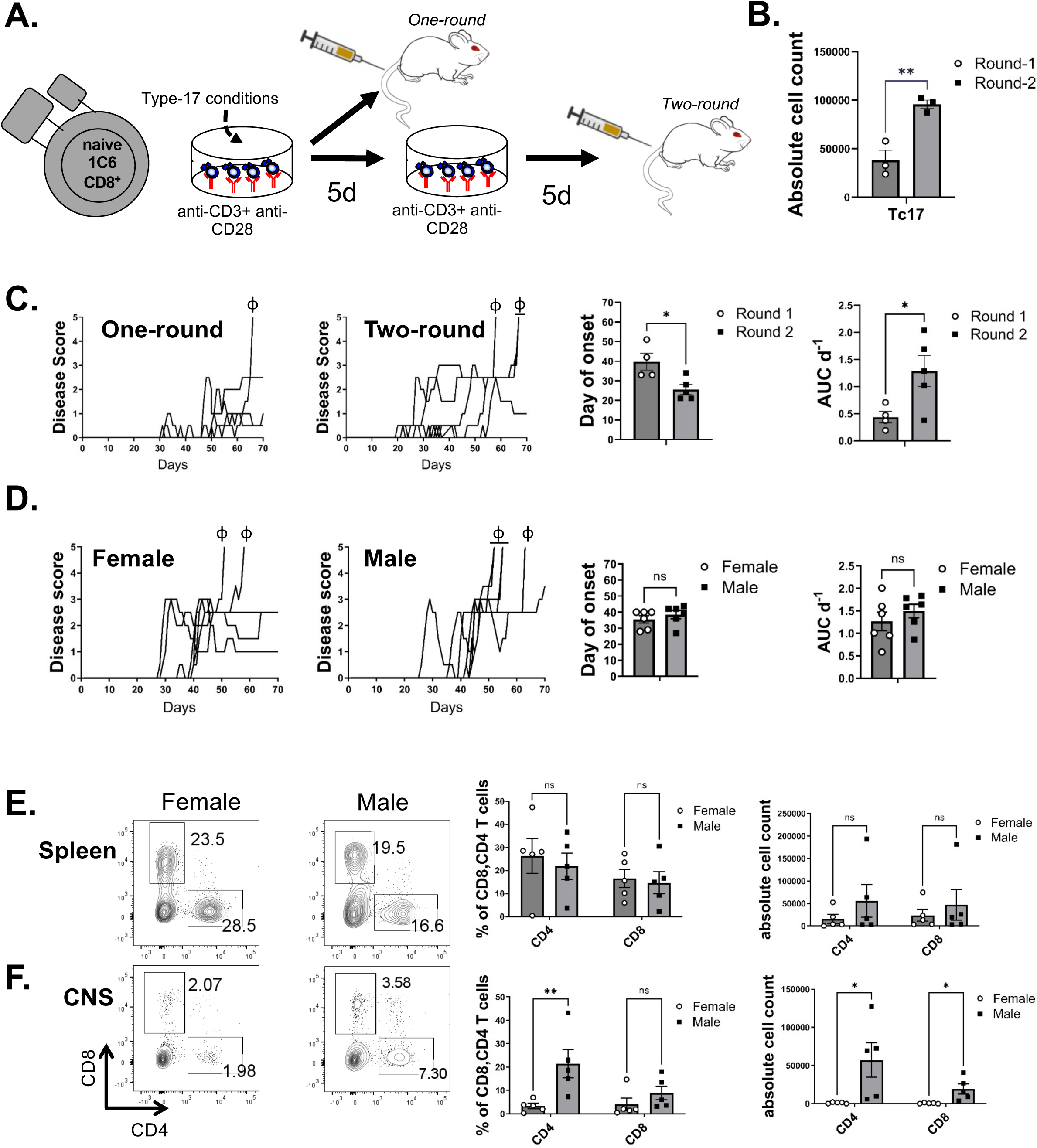
Male and female 1C6 Tc17 cells induce EAE of similar onset and severity. **A.** Schematic depicting experimental approach with restimulated Tc17 cells. **B.** Comparison of live CD8^+^ T cell yield from Tc17 cultures after one versus two rounds of stimulation. Representative of 2 experiments. **C.** Female Tc17 cells were generated using one-round and two-round stimulation protocols, were adoptively transferred to female NOD.*Scid* recipients and were monitored daily for signs of EAE. One-round recipients, n=4; two-round recipients, n=5. **D.** Tc17 cells were generated from female and male 1C6 CD8^+^ T cells using the two-round protocol and were adoptively transferred to sex-matched recipients (n=6, both groups) that were monitored daily for signs of EAE. **E, F.** Splenic (**E**) and CNS-infiltrating (**F**) T cells were isolated from recipient animals in Figure 5D and were assessed for expression of CD8 vs CD4. Each symbol represents an observation from a distinct mouse. ϕ, mouse attained ethical endpoint. *, p<0.05; **, p<0.01; n.s., not significant; two-tailed *t*-test. Related to Supplementary Figure 1.

We had previously shown that adoptive transfer of male 1C6 Th17 cells caused disease of higher overall severity relative to female Th17 counterparts; this was irrespective of the sex of the recipient mice (28). We thus addressed whether 1C6 Tc17 pathogenicity was similarly regulated by biological sex. Both female (3/6) and male (5/6) Tc17 cells could induce a progressive disease course in sex-matched recipients with similar onset and overall disease burden (Fig. 5D). Thus, unlike Th17 cells, 1C6 Tc17 cells did not display sex-driven differences in pathogenicity.

### Ex vivo analysis of T cells from Tc17 recipients

We next assessed T cell frequency and inflammatory markers from recipients of male versus female Tc17 cells at experimental endpoints. In spleens, the frequency and absolute number of CD8^+^ T cells was no different between spleens of female versus male Tc17 recipients (Fig 5E). Curiously, however, a substantial frequency of CD4^+^ T cells were observed in recipients of both male and female cells. This was despite having transferred Tc17 cells derived from cell-sorter-purified CD8^+^ T cell populations (Supp. Fig. 1A) and the fact that NOD.*Scid* mice lack endogenous lymphocytes.

To assess T cells within in the target organ, we quantified their presence in the CNS of mice that received male or female Tc17 cells. While the relative frequency of CD8^+^ T cells was no different in the CNS of recipients of male versus female cells, those mice that received male Tc17 cells showed a greater absolute number of CD8^+^ T cells (Fig 5F). Thus, male sex appeared to augment the capacity of effector Tc17 cells to infiltrate the CNS, despite having no apparent impact on disease burden. Notably, both the frequency and absolute number of CD4^+^ T cells were elevated in the CNS of recipients of male Tc17 cells (Fig 5F).

We then measured inflammatory cytokine expression from T cells isolated *ex vivo* from these adoptive transfer recipients. A substantial frequency of CD8^+^ T cells from both spleen and CNS were positive for IL-17, IFNγ and TNF, with no differences noted between the sexes (Supp. Fig 1B). CD4^+^ T cells were robustly positive for IFNγ and TNF but were less so for IL-17; again, no sex differences were observed (Supp. Fig. 1C). Next, assessment of the cytotoxic molecules Granzyme B and perforin revealed that a modest proportion of CD8^+^ (Supp. Fig 1D) and CD4^+^ (Supp. Fig 1E) T cells were positive for these markers, irrespective of sex or compartment.

### CD4^+^ T cells arise in NOD.Scid recipients of 1C6 Tc cells

We now wanted to further investigate our observation that CD4^+^ T cells are found in the spleen and CNS upon adoptive transfer of 1C6 Tc17 cells. We first showed that CD4^+^ T cells also arise in NOD.*Scid* recipients of 1C6 Tc1 cells and are present at similar frequencies as CD8^+^ T cells within the CNS (Supp. Fig. 1F). Thus, the accumulation of CD4^+^ T cells in vivo was observed irrespective of whether Tc1 or Tc17 cells were transferred. Notably, in this experiment, 1C6 CD8^+^ T cells were purified twice by cell sorting: first, prior to differentiation of CD8^+^CD62L^hi^ splenic T cells and again at the end of 5 days of Tc1 culture.

To study the kinetics of CD4^+^ and CD8^+^ T cell expansion in vivo, we adoptively transferred Tc17 cells to NOD.*Scid* recipients that we sacrificed 7, 14 or 21 days later. In both spleens and CNS, the overall frequency of T cells increased substantially over time. In spleens, a similar frequency of CD4^+^ and CD8^+^ T cells were noted at both day 7 and day 21, while in CNS, a similar frequency of CD4^+^ and CD8^+^ T cells were observed at all timepoints (Fig. 6A). We then adoptively transferred 1C6 Th17 cells to NOD.*Scid* mice and again assessed the kinetics of CD4^+^ and CD8^+^ T cells in spleen and CNS. No accumulation of CD8^+^ T cells were noted in these Th17 recipients (Fig. 6B). Thus, while CD4^+^ T cells were observed to arise in NOD.*Scid* mice upon adoptive transfer of 1C6 Tc1 or Tc17 cells, no comparable accumulation of CD8^+^ T cells was seen in recipients of 1C6 Th17 cells.

**Figure 6.**
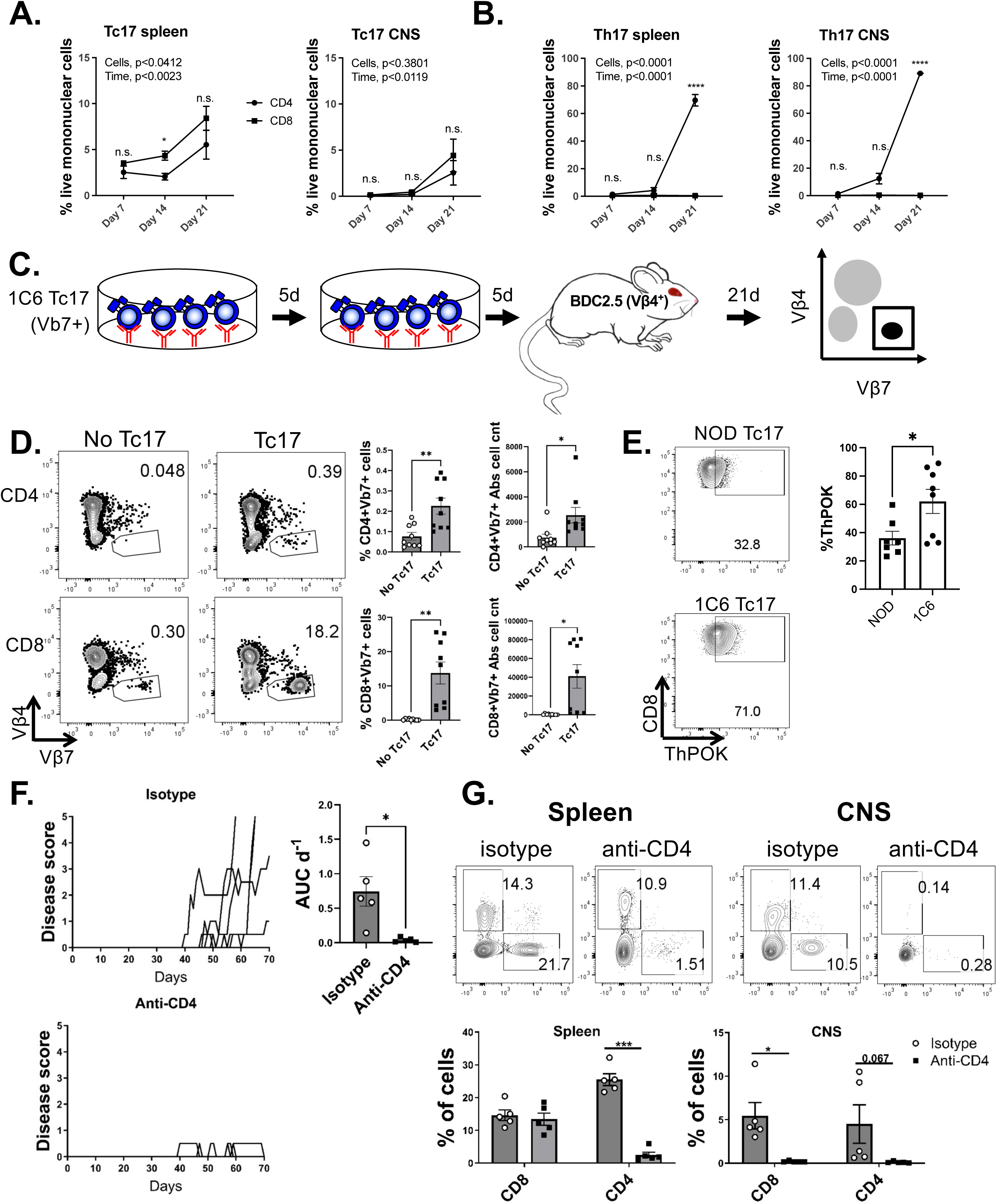
CD4^+^ T cells arise in 1C6 Tc17 recipients and are essential for disease. **A, B.** Female 1C6 Tc17 (**A**) or Th17 (**B**) cells were adoptively transferred to female NOD.*Scid* recipients (n=9 each). At each timepoint, n=3 mice were sacrificed and the presence of CD8^+^ vs CD4^+^ T cells was assessed from the indicated tissue compartment by flow cytometry. **C.** Schematic depicting the transfer of female 1C6 Tc17 cells to BDC2.5 recipients. **D.** Expression of Vβ7 vs Vβ4 on CD4^+^ and CD8^+^ splenic T cells from BDC2.5 mice that received, or did not receive, 1C6 Tc17 cells. The % frequency and absolute number of CD4^+^Vβ7^+^ and CD8^+^Vβ7^+^ T cells were quantified. Data are from two experiments with each symbol representing a unique recipient. **E.** Tc17 cells were differentiated over two rounds from female WT NOD or 1C6 CD8^+^ T cells and the expression of ThPOK was assessed. **F.** Female 1C6 Tc17 cells were differentiated using the two-round protocol and were adoptively transferred to two sets of female NOD.*Scid* recipients; one that received anti-CD4 antibody 2x weekly until d35, and another that received isotype control (both n=5). Recipients were monitored daily for signs of EAE. **G.** CD4^+^ and CD8^+^ T cells were assessed from the spleen and CNS of recipient animals at endpoints by flow cytometry. Quantification based on live CD8^+^ or CD4^+^ events. Each symbol represents an observation from a distinct mouse. *, p<0.05; **, p<0.01; ****, p<0.0001; n.s., not significant. Repeated measures ANOVA and Sidak’s multiple comparisons test (**A, B**); two-tailed *t*-test (**D-G**). Related to Supplementary Figure 1.

Despite having purified CD8^+^ T cells prior to transfer, it was still possible that we had also transferred a very small number of CD4^+^ T cells that subsequently underwent homeostatic expansion in lymphocyte-deficient NOD.*Scid* hosts. We therefore wanted to examine whether adoptive transfer of 1C6 Tc17 cells would give rise to CD4^+^ T cells in lymphocyte-sufficient mice with an intact CD4^+^ T cell compartment. We transferred Tc17 cells to NOD-background BDC2.5 mice in which >95% of T cells bear a transgenic Vβ4^+^ TcR (33) that can be readily distinguished from Vβ7^+^ 1C6 T cells. After 21 days, we assessed the presence of Vβ7^+^ T cells in the spleens of BDC2.5 mice that had received 1C6 Tc17 cells versus those that had not (Fig. 6C). We noted that both the frequency and absolute number of Vβ7^+^ CD4^+^ T cells were significantly increased in these animals relative to BDC2.5 mice that did not receive 1C6 Tc17 cells (Fig 6D) Unsurprisingly, the frequency and absolute number of Vβ7^+^ CD8^+^ T cells was higher in BDC2.5 mice that received 1C6 Tc17 cells.. Thus, CD4^+^ T cells arise from the transfer of 1C6 Tc17 cells to lymphocyte-sufficient mice.

CD4^+^ T cell lineage commitment is enforced in the thymus by the transcription factor ThPOK that concomitantly suppresses CD8 expression. Furthermore, its enforced expression is sufficient to redirect class-I restricted cells to the CD4 lineage (34). Interestingly, we found an increase in ThPOK expression in 1C6 Tc17 cells relative to those differentiated from WT NOD CD8^+^ T cells (Fig. 6E). Our data therefore indicated that 1C6 Tc17 cells express ThPOK, the master transcription factor for the CD4 lineage, and give rise to CD4^+^ T cells in the *in vivo* setting.

### CD4^+^ T cells are essential to 1C6 Tc17-mediated EAE

To determine whether Tc17-derived CD4^+^ T cells were essential to EAE, we transferred 1C6 Tc17 cells into NOD.*Scid* recipients followed by treatment with either anti-CD4 or isotype control. Strikingly, CD4 depletion sharply reduced disease severity in recipient mice, indicating that the CD4^+^ T cells that arise upon Tc17 transfer are essential to disease outcomes (Fig. 6F). We next examined the relative accumulation of CD8^+^ and CD4^+^ T cells in anti-CD4-treated and isotype control recipients. Anti-CD4 treatment reduced the frequency of CD8^+^ T cells that infiltrated the CNS, though the frequency of CD8^+^ T cells in the spleen did not change (Fig. 6G). Thus, CD4^+^ T cells are essential to Tc17-mediated disease induction and to CD8^+^ T cell infiltration of the target organ in our model.

## Discussion

Influenced in part by evidence from animal models (35), MS has traditionally been considered a CD4^+^ T cell-mediated disease. However, several lines of evidence point to CD8^+^ T cells as important players. CD8^+^ T cells outnumber CD4^+^ T cells in active lesions (10) and do so in progressive MS by as much as 50-fold (9). Further, the presence of injured axons in MS lesions correlates with the number of CD8^+^ T cells (36) and MHC class I alleles (A3 and B7) are linked to MS incidence (37). In chronic progressive MS lesions, CD8^+^ T cells display a tissue-resident effector memory cell phenotype, which show focally and temporally restricted activation (38). Intriguingly, the characteristic hallmark of numerous newly discovered autoimmune or paraneoplastic neurological diseases are inflammatory reactions that are dominated, and apparently mediated, by CD8^+^ T cells in the CNS (39, 40). Further, it has recently been proposed that CD8^+^ T cells in MS lesions are directed towards EBV-epitopes rather than towards myelin-derived antigens (41, 42). A better understanding of CD8^+^ T cell-driven pathogenic mechanisms in inflammatory CNS autoimmune disorders is therefore warranted.

There have been multiple reports of MHC class II-restricted CD8^+^ T cells in the context of chronic viral infection. Such cells are present in a small proportion of HIV-1-infected elite controllers (43) and expand in rhesus macaques vaccinated with simian immunodeficiency virus (SIV) antigen-expressing cytomegalovirus (44). However, a possible role for such cells in autoimmunity is less well appreciated. In rodents, class II-restricted CD8^+^ T cells have been reported, but in the context of *Cd4^-/-^* mice (45–47) in which unconventional CD8^+^ T cells may arise to compensate for the lack of CD4^+^ T cells. As 1C6 mice unexpectedly possess CD8^+^ T cells that recognize MOG_[35-55]_ cognate antigen in an MHC class II-dependent manner (27), this afforded us the opportunity to study the function of these unusual cells in CNS autoimmunity.

It had previously been shown that 1C6 CD8^+^ T cells proliferate in response to MOG_[35-55]_ in a class II-dependent manner. However, a head-to-head comparison of 1C6 CD8^+^ to CD4^+^ T cells revealed that the former displayed ∼3-4 fold lower [^3^H]-thymidine incorporation upon MOG_[35-55]_ stimulation (27). Here, we found that while 1C6 Tc1 and Tc17 cultures expanded less robustly than Th1 and Th17 comparators, the cell-intrinsic proliferative capacity of MOG_[35-55]_ 1C6 Tc cells was unimpaired. Further, 1C6 Tc1 cells capably produced the signature cytokine IFNγ when cultured with MOG_[35-55]_, while Tc17 cells produced both IFNγ and IL-17. Thus, we confirmed that 1C6 CD8^+^ Tc cells respond to the cognate antigen MOG_[35-55]_.

Several models of CNS antigen-specific CD8^+^ T cell-mediated EAE have been previously described. Adoptive transfer of myelin basic protein (MBP)-reactive CD8^+^ T cell clones caused atypical EAE characterized clinically by ataxia (18) while mice bearing a class I-restricted transgenic TcR with specificity for the astrocytic marker glial fibrillary acidic protein (GFAP) developed spontaneous ataxic disease (48). As we recently demonstrated that adoptive transfer of 1C6 Th1 or Th17 cells elicited chronically worsening CNS autoimmunity in recipient mice (28) and as 1C6 mice possess transgenic CD8^+^ T cells, we asked whether transfer of 1C6 Tc1 and Tc17 cells induced similar disease. Indeed, both Tc1 and Tc17 1C6 cells transferred disease with a similar severity as seen in recipients of Th1 and Th17 1C6 cells, albeit with a delayed onset. Interestingly, we noted a large thalamic lesion with in a Tc1 recipient mouse. Focal thalamic lesions characterize most cases of MS (31, 32, 49). Further work is required to fully understand a potential contribution of effector 1C6 CD8^+^ T cells to deep grey matter injury.

Surprisingly, we noted a robust presence of CD4^+^ T cells in the spleens and CNS of NOD.*Scid* mice that received 1C6 Tc effector T cells. These CD4^+^ T cells were essential to Tc-mediated disease and to the infiltration of CD8^+^ T cells into the CNS. Our findings are concordant with those of Leuenberger and colleagues (50) who, using a model in which *Rag1^-/-^* mice were reconstituted with CD8^+^ T cells alone and then immunized with MOG_[35-55]_, found that CD4^+^ T cells arise in recipients that show clinical signs of EAE. Importantly, we also provide evidence that 1C6 Tc17 cells can give rise to CD4^+^ T cells in immunocompetent hosts. An elegant recent study (51) used two different class I-restricted TcR transgenic mouse lines to show that CD8^+^ T cells can cross-differentiate into CD4^+^ T cells in the in vivo setting. This property was unrelated to self/non-self-specificity, as one strain (OT-I) recognized exogenous antigen, while the other (8.3) was specific for a pancreatic self-Ag. Cross-differentiated CD4^+^ T cells were observed at the highest frequency in the gut lamina propria (LP) and mesenteric lymph nodes, and it was posited that gut microbiota are responsible for the expansion of these cells. Nevertheless, cross-differentiated OT-I CD4^+^ T cells were also seen at detectable frequencies (>1%) in the spleen. Thus, this property is not unique to class II-restricted CD8^+^ T cells such as is the case in 1C6 mice (27). Furthermore, transfer of mature polyclonal CD8^+^ T cells also caused the expansion of a robust population of CD4^+^ T cells (>20%) in mesenteric lymph nodes (51), suggesting that the capacity of CD8^+^ T cells to transdifferentiate into CD4^+^ T cells is not exclusive to TcR-transgenic lines in which the T cell repertoire is highly skewed. In the future, it would be interesting to further explore the molecular mechanisms that underpin the capacity of 1C6 T cells to give rise to CD4^+^ T cells in vivo, building on our observation that 1C6 CD8^+^ T cells show elevated expression of the CD4 lineage determining factor ThPOK.

While CD4^+^ T cells are essential to 1C6 Tc17-mediated EAE, we observe no differences in disease severity between recipients of male vs female Tc17 cells: this contrasts with our previous observation that male 1C6 Th17 cells induce disease of greater severity than female counterparts. These data argue against the possibility that the CD4^+^ T cells that arise upon Tc17 transfer are identical to bonafide Th17 cells. Intriguingly, 1C6 Tc17 cells are absent from the CNS upon anti-CD4 treatment. This raises the possibility that the CD4^+^ T cells seen in Tc17 recipients contribute to pathogenicity by facilitating Tc17 entry into the CNS: indeed, Leuenburger and colleagues have seen that CD8^+^ T cell motility within the CNS requires the presence of CD4^+^ T cells (50). Further work is needed to characterize the phenotype and contributions of CD4^+^ T cells to EAE in CD8^+^ T cell recipients.

In conclusion, adoptive transfer of myelin antigen-specific effector CD8^+^ T cells can induce EAE with a chronically worsening clinical course. Intriguingly, Tc cell transfer gives rise to CD4^+^ T cells that are essential to for both disease and for CD8^+^ T cell infiltration of the CNS. We therefore describe a new model with which to study the collaboration of CD4^+^ and CD8^+^ T cells in the pathogenesis of chronic progressive CNS autoimmunity.

## Acknowledgments

We thank Vijay K. Kuchroo for providing us with the 1C6 line, Kim Larose-Labrecque and Cindy Ouellet for animal care and Vincent Desrosiers for assistance with flow cytometry.

## Materials and Methods

### Experimental animals and ethics

1C6 mice have been previously described (27) and were maintained in our animal facility at the CHU de Québec. NOD.Cg*-Prkdc^scid^*/J mice (#001303; NOD.*Scid*) were obtained from the Jackson Laboratory and maintained in our animal facility. NOD.Cg-Tg(TcraBDC2.5, TcrbBDC2.5)1Doi/DoiJ mice (#004460, BDC2.5) were obtained from the Jackson Laboratory. Mice were co-housed. All animal breedings and experiments were approved by the Animal Protection Committee of Université Laval (protocols 2023-1375, 2023-1376, 2021-830).

### Effector Tc and Th culture and differentiation

CD8^+^ and CD4^+^ T cells were isolated from 1C6 spleens and LNs using anti-CD8 and anti-CD4 MicroBeads (Miltenyi), respectively. Naïve CD8^+^CD62L^hi^ and CD4^+^CD62L^hi^ T cells were purified using a FACSAria (BD) high-speed cell sorter, and were cultured in supplemented T cell media as described (53). For plate-bound stimulation, T cells were differentiated for 2 days on plates coated with anti-CD3 and anti-CD28 (4 µg mL^-1^ for CD8 and 2ug mL^-1^ for CD4; BioXcell) and were subsequently transferred to uncoated plates for an additional 3 days. For stimulation with cognate peptide+APCs, T cells were cultured for 5 days at a 1:5 ratio with irradiated splenocytes (2000 cGy) pulsed with 2 μg mL^-1^ of MOG_[35-55]_ (CHU de Québec). For cell proliferation analysis, T cells were incubated with CellTrace Violet (Fisher Scientific) prior to stimulation. For type-1 (Tc1/Th1) differentiation, T cells were cultured with 10 ng mL^-1^ rmIL-12 (R&D Biosystems) plus anti-IL-4 (10 µg mL^-1^, BioXcell) for 2 days, and with 10 ng mL^-1^ rmIL-2 (Miltenyi) for another 3 days. For type-17 (Tc17/Th17) differentiation, T cells were cultured for 2 days with 3 ng mL^-1^ rhTGFb (Miltenyi) + 20 ng mL^-1^ rmIL-6 (Miltenyi) + 10 µg mL-1 anti-IFNγ (BioXcell), and with 20 ng mL^-1^ rmIL-23 (R&D Biosystems) for an additional 3 days (30).

### Flow cytometry

Cells were pre-incubated with Fc Block (BD Biosciences) to prevent non-specific antibody binding and then stained with cell surface markers: CD8 (clone 53.6.7 ThermoFisher), CD4 (clone RM4.5, ThermoFisher), Vβ7 (clone TR310, Biolegend), Vβ4 (clone KT4, BD Biosciences) for 20 min at 4⁰C. Cells were also stained with Fixable Viability Dye (ThermoFisher) for 15min; positive (dead) cells were gated out in all analyses. We enumerated absolute cell counts with 123count eBeads (ThermoFisher). For intracellular cytokine staining, cells were first incubated for 4 hrs in the presence of 50 ng mL^-1^ phorbol 12-myristate 13-acetate (Sigma-Aldrich), 1µM ionomycin (Sigma-Aldrich) and Golgi Stop (1 µL per mL culture; BD Biosciences) and then stained for surface antigens and live/dead indicator. Cells were processed using Fixation and Perm/Wash buffers (eBioscience) and were stained for intracellular antigens using the following antibodies: TNF (clone MP6-XT22 ThermoFisher), IFNγ (clone XMG1.2, ThermoFisher), IL-17A (clone TC11-18H10.1, ThermoFisher), Granzyme B (clone NGZB, ThermoFisher), Perforin (clone eBio0MAK-D, ThermoFisher). Flow cytometry data were collected by using FACS Canto II flow cytometer (BD Biosciences) and were analyzed using FlowJo software. Gates were set on the basis of fluorescence minus one controls.

### Adoptive transfer EAE

1C6 Tc1, Tc17, Th1 or Th17 cells were generated by plate-bound stimulation as described above and were administered i.v. to NOD.*Scid* mice (5×10^6^ cells per mouse). For Th+Tc co-transfers, 2.5×10^6^ of each cell type were transferred at the same time. In some experiments, Tc cells were restimulated for a second 5-day round of differentiation as indicated in the Figure legends. Recipient animals were treated i.p. with 200 ng pertussis toxin on day 0 and day 2. They were weighed and scored daily up to 70 days using a well-established semiquantitative scale that we have used previously (54): 0, no disease; 1, loss of tail tone; 2, hind limb weakness or partial paralysis; 3, complete hind limb paralysis; 4, front and hind limb paralysis; 5, ethical endpoints attained. Mice with advanced symptoms (score ≥3) were monitored daily by trained animal technicians, in collaboration with the veterinary service of Université Laval, to assess whether ethical endpoints had been attained. Individual disease curves were assessed for day of symptom onset, disease burden (area under curve) and for progression. A mouse was considered to have developed disease if it attained a minimum score of 1. Progression was evaluated based on our described paradigm (28). Briefly, a course was considered “progressive” if it met four criteria: *i)* an increase in disease severity of ≥ 1 over 10 days, *ii)* did not have a single-day severity decrease of > 0.5 during the same 10 day period, *iii)* did not remit after the 10 day period, *iv)* exceeded a severity score of 2.5 during the 10 days or thereafter.

### Histopathology

Mice were euthanized and perfused with cold PBS followed by 4% paraformaldehyde (PFA) via the left cardiac ventricle. The brains and spinal cords were carefully extracted from the skull and spinal column, respectively. Tissues were then stored in 4% PFA at 4°C for 24 hours, followed by at least 48 hours in PBS before being embedded in paraffin. Thin sections (5 μm) of the brain, and spinal cord were stained with hematoxylin & eosin (H&E), Luxol fast blue for myelin, or Bielschowsky’s silver impregnation for neurons and axons (55).

### CNS mononuclear cell isolation

Mice were euthanized and perfused with cold PBS administered through the left cardiac ventricle. CNS was dissected from the skull and spinal column, respectively. CNS tissue was homogenized using a PTFE Tissue Grinder (VWR) and was incubated at 37⁰c for 30 min in homogenization solution (HBSS containing 4 ng mL-1 liberase and 25 ng mL-1 DNase). Homogenate was filtered through a 70-µm cell strainer, resuspended in 35% Percoll (GE Healthcare) and centrifuged. Mononuclear cells were collected, washed and prepared for flow cytometric analysis.

### CD4 blockade

Starting from day 0 of adoptive transfer of Tc17 cells, CD4^+^ T cells were depleted by treating the mice with intraperitoneal injection of 400ug/mouse of anti-CD4 (clone GK1.5, BioXCell) in PBS, twice weekly to day 35. Anti-keyhole limpet hemocyanin (clone LTF-2, BioXCell) was used as an isotype control.

### Statistics

Statistical analyses were conducted using Prism software (GraphPad). Information regarding the specific tests used are indicated in individual Figure legends. Two-tailed tests were used in each case. Comparisons between multiple groups or timepoints were conducted using ANOVA followed by a post-hoc test.

---

**Supplementary Figure 1. Data related to Main Figure 5. A.** Representative pre- and post-sort flow cytometry plots of female peripheral CD8^+^ T cells that were enriched using anti-CD8-labeled magnetic beads (*pre-sort*) and subsequent purification of CD8^+^CD4^-^ cells by high-speed cell sorting (*post-sort*). Sorter-purifed CD8^+^ T cells were then used for Tc17 cultures. **B, C.** CD8^+^ (**B**) and CD4^+^ (**C**) T cells from recipient mice in Figure 5D were assessed for expression of the indicated cytokines. **D, E.** CD8^+^ (**D**) and CD4^+^ (**E**) T cells from recipient mice were assessed for expression of GzB and Pzf. **F.** Male Tc1 cells were generated from sorted CD8^+^CD62L^hi^ splenic 1C6 T cells, were again purified on CD8 positivity after 5 days’ culture and were adoptively transferred to male NOD.*Scid* mice. At experimental endpoints, splenic and CNS mononuclear cells were assessed for positivity of CD4 and CD8. **, p<0.01; n.s., not significant; *t*-test. Each data point represents an individual mouse.

**Supplementary Table 1. Histopathological characteristics of Tc recipient mice.** Representative image from mouse 3 is presented in main Figure 4A. Representative image from mouse 1 is presented in main Figure 4B.

