## Supplementary material for "Myelin antigen-specific effector CD8^+^ T cells induce chronic CNS autoimmunity in a CD4^+^ T cell-dependent manner": Supp Fig 1 Supp Table 1

### Supplementary Figure 1

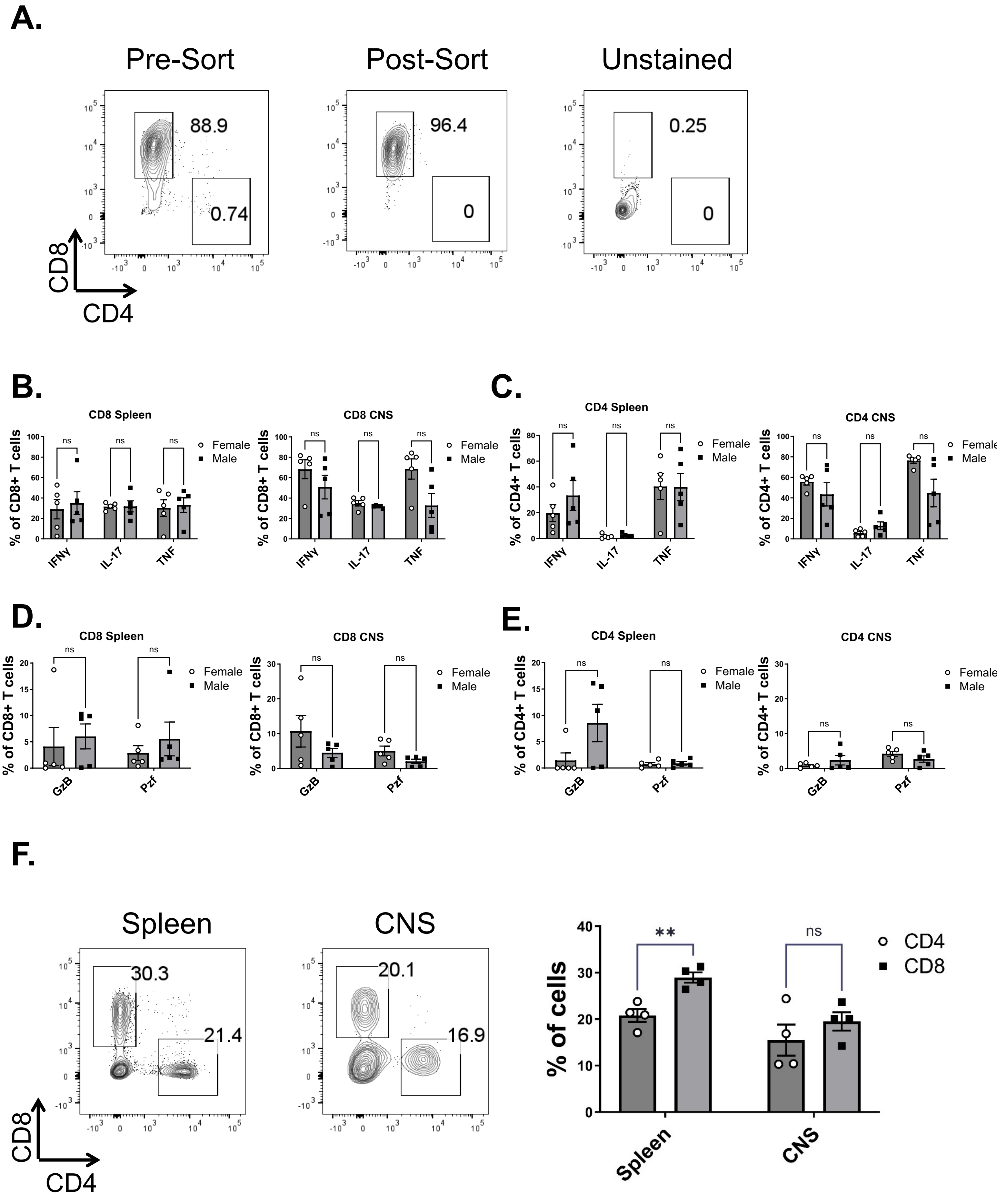

### Supplementary Table 1

| | Cells transfered | Spinal cord | Active lesions? | Brain | Active lesions? |
|---|---|---|---|---|---|
| 1 | Female Tc1 | Large confluent demyelinating lesions, lateral and dorsal column | Mainly active | Large confluent demyelinating lesions; thalamus, mesencephalon, medulla | active and inactive |
| 2 | Female Tc1 | Mild meningeal inflammation | - | Profound inflammation; small to moderately sized inflammatory demyelinating lesions; fornix, medulla | active |
| 3 | Female Tc1 | Large confluent demyelinating lesions, anterior column; signs of early Schwann cell remyelination | active | 3 moderately sized inflammatory demyelinating lesions in fornix, mesencephalon; signs of Schwann cell demyelination | mainly active |
| 4 | Female Tc17 | Single inflammatory demyelinating lesion | inactive | none | - |
